# Uncovering the mosaic evolution of the carnivoran skeletal system

**DOI:** 10.1101/2023.09.03.556127

**Authors:** Chris J. Law, Leslea J. Hlusko, Z. Jack Tseng

## Abstract

The diversity of vertebrate skeletons is often attributed to adaptations to distinct ecological factors such as diet, locomotion, and sensory environment. Although the adaptive evolution of skull, appendicular skeleton, and vertebral column is well studied in vertebrates, comprehensive investigations of all skeletal components simultaneously are rarely performed. Consequently, we know little of how modes of evolution differ among skeletal components. Here, we tested if ecological and phylogenetic effects led to distinct modes of evolution among the cranial, appendicular, and vertebral regions in extant carnivoran skeletons. Using multivariate evolutionary models, we found mosaic evolution in which only the mandible, hindlimb, and posterior (i.e., last thoracic and lumbar) vertebrae showed evidence of adaptation towards ecological regimes whereas the remaining skeletal components reflect clade-specific evolutionary shifts. We hypothesize that the decoupled evolution of individual skeletal components may have led to the origination of distinct adaptive zones and morphologies among extant carnivoran families that reflect phylogenetic hierarchies. Overall, our work highlights the importance of examining multiple skeletal components simultaneously in ecomorphological analyses. Ongoing work integrating the fossil and paleoenvironmental record will further clarify deep-time drivers that govern carnivoran diversity we see today and reveal the complexity of evolutionary processes in multicomponent systems.

## Introduction

The diversity of animal forms is one of the most salient patterns across the tree of life. In mammals, morphological innovations in the skull, appendicular skeleton, and vertebral column facilitate the incredible diversity found today, ranging from bats with winged forelimbs to the biggest animals to have ever lived on earth. Many researchers have examined how variation in the skull [1–7], limbs [8–11], or vertebrae [12–16] serve as crucial adaptations to their evolution. These skeletal systems are traditionally examined independently and are rarely investigated simultaneously even though these anatomical regions comprise a single, functionally integrated system that serves as structural support for movement, locomotion, and other life functions. When considered wholistically, the observed variation across the different components of organismal anatomy is generally explained by multitudinous factors, some that are potentially incongruous [17–20]. While this evolutionary push-and-pull between anatomical regions may characterize the process of evolution, the hypothesis can only be tested when the different skeletal components are explored simultaneously rather than piecemeal. Simultaneous investigation of integrated components is critical to our understanding of the role of developmental and/or functional integration in canalizing macroevolutionary trajectories [21– 23]. Here, we use carnivorans to investigate how ecological and phylogenetic factors correspond to evolutionary changes in the cranial, appendicular, and axial skeletal systems. Carnivorans (bears, cats, dogs, seals, and their relatives) are a productive model system to examine skeletal evolution because of their high species richness and vast distribution across most biomes in all continents and oceans, along with broad ecological diversity in locomotor traits and feeding adaptations.

Components of carnivoran skeletal systems are well studied individually. In the skull, craniomandibular diversity is influenced by several ecological factors and phylogeny [24–28]. The skull exhibits decoupled evolutionary modes: cranial shape follows clade-specific evolutionary shifts, whereas mandibular shape evolution is linked to broad dietary regimes [6,29]. In the appendicular skeleton, ecomorphological divergence exists between the hindlimbs, which are adapted primarily for locomotion, and the forelimb, which are adapted for multiple functions ranging from running to grappling prey to manipulating objects [30–33]. Additionally, more recent work using phylogenetic comparative methods found that scaling and phylogeny exhibit stronger effects on limb evolution than do ecological parameters [34–36]. In contrast to craniomandibular and appendicular ecomorphology, research on the axial skeleton is in nascent stages. Initial research indicates that distinct regions of the vertebral column are under different evolutionary pressures. Anterior (i.e., more cranial) vertebrae exhibits low disparity due to phylogenetic constraints or ecological conservatism, whereas posterior (i.e., more caudal) vertebrae exhibits higher disparity that may be due to adaptations to various locomotor ecologies [13,37]. In contrast to these morphologically-localized studies, analyses of the evolution of whole-body traits like body mass, skeletal size, and body shape often follow a Brownian motion model or clade-based shift model rather than being associated with ecological regimes [28,38,39].

Compared to skeletal system-specific findings, simultaneous investigation of skulls, limbs, vertebrae, and overall body plan are rarely conducted, likely because of the enormous amount of data that would need to be collected and the complexity of the multivariate analyses required. However, a more comprehensive approach to quantifying skeletal evolution is essential to elucidate its complexity more fully. The search for system-level trends and variations is further obscured by the disparate methods employed to test the effects of ecology and phylogeny on different skeletal systems by different researchers. In this study, we address both issues in our investigation of the mosaic evolution of carnivoran skeletons by creating a new phenomic dataset that encompasses all major components of the skeletal system and using a unified set of multivariate evolutionary models to test the ecological and phylogenetic effects influencing the modes of evolution of these skeletal components.

## Methods

### Skeletal and ecological traits

We collected 103 linear measurements to capture the skeletal morphology of 119 carnivoran species (208 osteological specimens; Fig. S1; Table S1). This dataset includes seven cranial traits, seven mandibular traits, 13 forelimb traits, 13 hindlimb traits, and seven traits in third cervical, fifth cervical, first thoracic, middle thoracic, diaphragmatic thoracic, last thoracic, first lumbar, middle lumbar, and last lumbar vertebrae. Because carnivorans exhibit differing degrees of sexual dimorphism [40,41], we use only male specimens. To remove size effects, we calculated log shape ratios by dividing each skeletal trait by the geometric mean of all 103 traits [42,43]. We then used principal component analyses (PCAs) to reduce the dimension of each skeletal component (i.e., cranium, mandible, forelimb, hindlimb, and each of the nine vertebrae) and retained a number of PC axes that corresponded to >90% of the explained variance. We also conducted a PCA on the entire dataset as our proxy of the whole-skeleton phenome and retained the first six PC axes (∼75% of explained variance) for subsequent analyses. We classified the 119 carnivoran species into distinct locomotor modes, hunting behaviors, and dietary regimes following [39].

### Phylogenetic comparative methods

We tested whether each skeletal component evolved as adaptation to specific ecological regimes or exhibited clade-specific evolutionary shifts by fitting multivariate evolutionary models on the retained PC axes of each skeletal component [44–46]. For the adaptive ecological models, we fit three multivariate multi-optima Ornstein-Uhlenbeck models (i.e., mvOUM_diet_, mvOUM_hunting_, and mvOUM_locomotion_) to test if dietary, hunting behavioral, or locomotor regimes influenced the evolution of each skeletal component using mvMORPH [46]. The models were fit across 500 stochastically mapped trees to account for uncertainty in phylogenetic topology and ancestral character states (see electronic supplementary materials). We also calculated the phylogenetic half-lives of the best supported adaptive ecological model [44]. A short phylogenetic half-life relative to the age of Carnivora (48.2 myr) would suggest that skeletal traits are strongly pulled toward distinct ecological optima across the adaptive landscape. For the clade-based model, we fit a multi-optima OU model (mvOUM_phyloEM_) without *a priori* ecological regimes with PhylogeneticEM [47]. We also fit a single-rate multivariate Brownian motion model (mvBM1) and a single-optimum OU model (mvOU1). We assessed the relative support of models using small sample-corrected Akaike weights (AICcW). Lastly, we assessed the covariation among skeletal components using partial least squares with geomorph [48].

Preliminary results revealed that phenotypic differences between pinnipeds (i.e., seals and sea lions) and terrestrial carnivorans are often the greatest source of variation for most skeletal components. These results are unsurprising considering pinnipeds exhibit derived morphologies that enable them to be fully aquatic. Therefore, we repeated our analyses using a reduced dataset with no pinnipeds. Results of the full dataset with pinnipeds are presented in the electronic supplementary material.

## Results and Discussion

We found mosaic evolution of the carnivoran skeleton in which ecology and phylogeny have differing influences on the evolutionary mode of the various skeletal components. Consistent with [6,29], the cranium and mandible exhibited decoupled evolutionary modes. In the cranium, the clade-specific shift model exhibited overwhelmingly greater support (mvOUM_phyloEM_; AICcW>0.99) compared to adaptive ecological models (Fig. 1; Table S2). We found eight evolutionary shifts in cranial morphology that correspond to carnivoran clades (Fig. 2A). In contrast, the adaptive dietary model was the best supported model (mvOUM_diet_; AICcW=0.96) for the mandible with a short phylogenetic half-life of 2.52 myr (Fig. 1; Fig. S2B; Table S2; see Supplementary Results for optima distribution in phylomorphospace). These results are congruent with findings revealing that mandibular shape is evolutionarily labile with respect to dietary evolution whereas cranial shape is partitioned among families rather than among dietary groups [6]. Despite their covariation (r = 0.73; Table S3), decoupled evolutionary modes between the cranium and mandible may be explained by their functions. Diet is often found to have had a strong influence on mandibular evolution because of its direct role in feeding [3,49–53]. In contrast, the cranium has multiple sensory functions in addition to feeding that influence its evolution [54–56], and therefore, the signal from dietary adaptations in its morphology may be obscured.

**Fig. 1.**
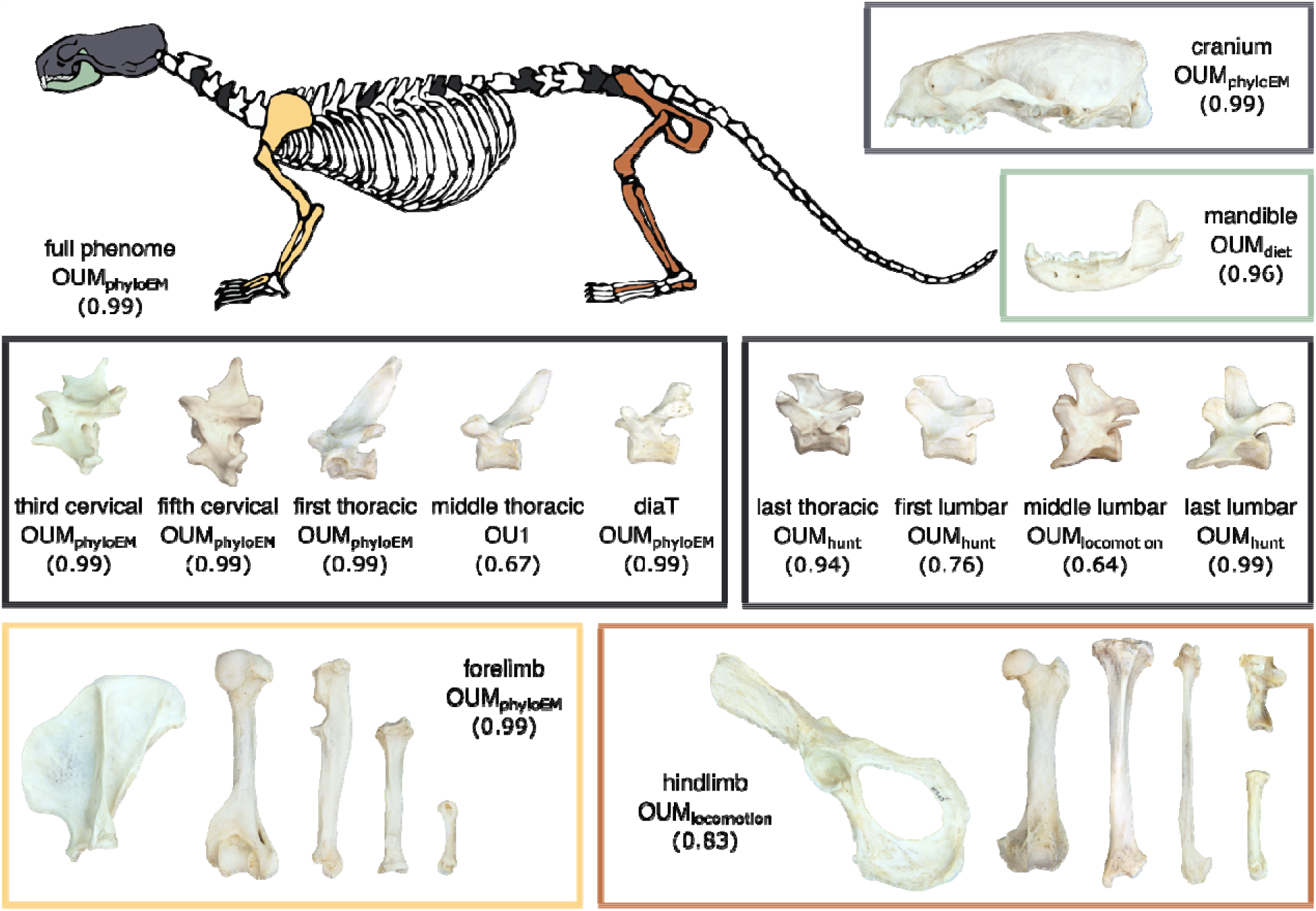
Diagram of the skeletal components and their best-fitting evolutionary model on *Lontra canadensis*. AICcW are in parentheses. See Table S2 for full AICc table. diaT = diaphragmatic thoracic vertebrae.

**Fig. 2.**
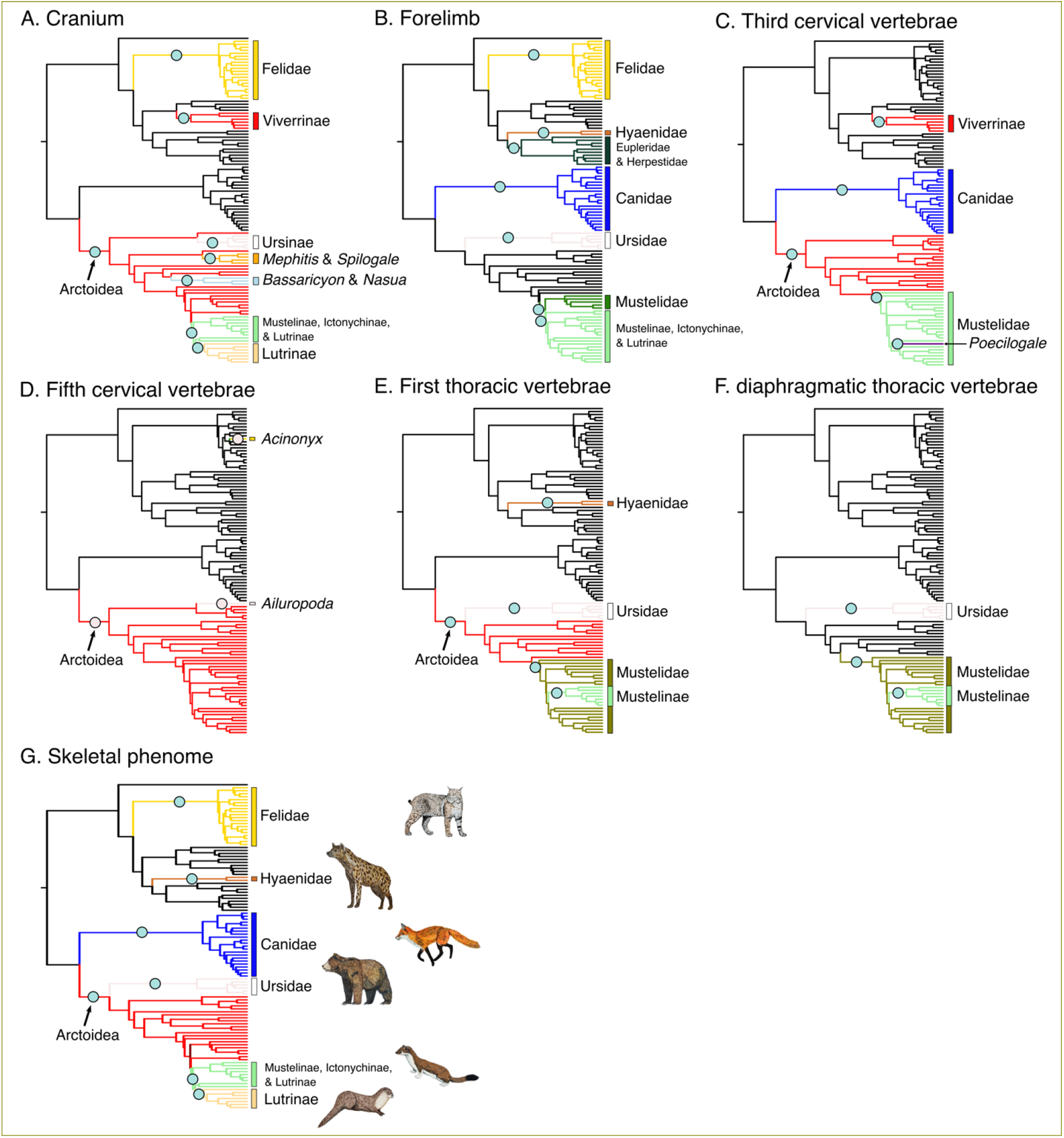
Clade-specific evolutionary shifts in skeletal components across terrestrial carnivorans identified by PhylogeneticEM. Shifts are represented as pink circles, and branches on the phylogenies are colored according to each regime.

The appendicular system exhibited decoupled evolutionary modes between forelimbs and hindlimbs. The forelimb was best supported by the mvOUM_phyloEM_ model (AICcW>0.99; Fig. 1; Table S2). Seven shifts in forelimb evolution occur primarily along familial branches (Fig. 2B), indicating that the complexity and variation of carnivoran forelimb morphology cannot be captured effectively by dietary, hunting behavioral, or locomotor categories. Instead, these shifts suggest that clade-specific adaptations enabled the diversity of forelimb skeletons for tasks such as grappling or manipulating prey, swimming, or digging [30–33,36,57,58]. For example, most felids use their prehensile forelimbs to ambush and subdue prey, most canids and hyaenids pounce and pursue prey, and some mustelids use their powerful forelimbs to dig out prey while other more derived mustelids (i.e., weasels) pursue prey in tight crevices and burrows [59]. In contrast, the hindlimb was best supported by the mvOUM_locomotion_ model (AICcW=0.83) in the hindlimb with a short phylogenetic half-life of 5.05 myr (Fig. 1; Table S2), supporting hypotheses that the hindlimb is adapted primarily for locomotion as typically found in quadrupedal mammals [60]. Although the forelimb and hindlimb covaries (r = 0.87; Table S3), previous work found that this integration is weaker than expected in carnivorans that do not specialize in cursoriality [36]. This work together supports the hypothesis of functional divergence between the forelimbs and hindlimbs of carnivorans.

The axial skeleton exhibits distinct evolutionary modes between the anterior and posterior regions of the vertebral column: cervical and most thoracic vertebrae tended to be best supported by clade-specific shift or single-peak OU models, whereas the last thoracic and all lumbar vertebrae were best supported by mvOUM_hunting_ or mvOUM_locomotion_ models (Fig. 1; Fig. 2C–F; Table S2). Our findings strengthen the coalescing hypothesis that anterior vertebrae exhibit lower disparity, higher evolutionary constraints, and more subtle adaptations to locomotion whereas posterior vertebrae exhibit the opposite patterns in carnivorans [37] and broadly across mammals [14]. We posit that high evolutionary constraints of the anterior vertebrae are associated with clade-specific shifts in the cervical and most thoracic vertebrae. Importantly, subtle adaptations in these anterior vertebrae could be masked by many-to-one or one-to-many mappings, making it difficult to uncover the form-function associations with evolutionary models [61]. In contrast, relaxed evolutionary constraints of the posterior vertebrae facilitate the evolution of disparate lumbar vertebrae across the entire carnivoran order. These disparate vertebrae adapt to diverse locomotor modes or hunting behaviors based on the mobility of the posterior vertebrae and irrespective of clade origins. The short phylogenetic half-lives (1.47–5.12 myr) further suggests strong pulls towards these different adaptive optima. More broadly, this increased mobility of the lumbar region over evolutionary time is hypothesized to be an innovation characterizing crown mammals [14,62,63]. Correspondingly, the posterior vertebrae are tightly integrated (r = 0.84–0.96; Table S3).

Lastly, we found that the clade-specific shift model (mvOUM_phyloEM_; AICcW > 0.99) best described the overall skeletal phenome (Table S2), a pattern that is consistent with previous investigations of whole-body proxies such as body size and body shape [28,38,39]. The mammalian body plan is comprised of cranial, axial, and appendicular components; therefore, its multidimensionality transcends one-to-one mapping relationships between morphology and ecological function. Instead, individual skeletal components within distinct body plans can adapt to specific ecological factors independently from each other, enabling species with distinct body plans to exhibit similar ecological or functional regimes and vice versa.

Overall, we elucidate the mosaic evolution of the carnivoran skeleton, finding that different skeletal components exhibit distinct modes of evolution. Our results suggest that different methodologies and taxonomic samples do not necessarily explain previously reported region-specific macroevolutionary patterns; rather, complexity in explanatory factors of skeletal diversity is a key feature of Carnivora. The ability of individual skeletal components to adapt to specific ecological factors independently from each other may have contributed to the clade’s *hierarchical* [64,65] evolution. As previously hypothesized [28,38], the restriction of carnassial shear to the P4/m1 pair may have been the key innovation that facilitated the initial carnivoran diversification early in the clade’s evolutionary history. Subsequent evolution led to the continual partitioning between clades, resulting in the origination of extant carnivoran families as discrete phylogenetic clusters that occupy different adaptive zones [66] with distinct morphologies including body size and shape [39,67] and various components of the skeleton ([6]; Fig. 2). Within-clade variation then arises to reflect resource partitioning among ecologically similar taxa, leading to adaptations in morphologies such as the mandible, hindlimb, and posterior region of the vertebral column (Fig. 1). These traits were strongly pulled toward distinct ecological peaks across the adaptive landscape as revealed by their short phylogenetic half-lives (1.47–5.12 myr) relative to the clade’s age (48.2 myr).

Our research statistically revealed the mosaic evolution of carnivoran skeletons. These distinct evolutionary modes demonstrate the importance of examining multiple skeletal components in ecomorphological analyses. Nevertheless, key questions remain: What spurred the evolutionary transitions towards the evolutionary shifts or adaptations of the various skeletal components? When in the 55 million years of carnivoran evolutionary history did these evolutionary events occur? And what developmental and genetic phenomena underlie the evolutionary dissociation of various skeletal elements? Ongoing work integrating the fossil and paleoenvironmental record will further elucidate the carnivoran diversity we see today and reveal the complexity of evolutionary processes in multicomponent systems.

## Acknowledgements

We are grateful to the staff and collections at the American Museum of Natural History, California Academy of Sciences, Field Museum of Natural History, Natural History Museum of Los Angeles County, Museum of Vertebrate Zoology, Natural History Museum London, San Diego Natural History Museum, Texas Vertebrate Paleontology Collection, National Museum of Natural History, and Burke Museum of Natural History and Culture. We thank Vera Weisbecker and 3 anonymous reviewers for their feedback.

## Data Accessibility Statement

All data and original code are made available on dryad (doi:10.5061/dryad.c2fqz61gf) [68].

