## Supplementary Materials for "Uncovering the mosaic evolution of the carnivoran skeletal system"

Electronic supplementary material for

Supplementary Methods

Supplementary Results and Discussion

Supplementary Figures

Figure S1. Skeletal trait measurements used in this study.

Figure S2. Phylomorphospaces of skeletal components and full phenome across terrestrial carnivorans (reduced dataset).

Figure S3. Diagram of the skeletal components and their best-fitting evolutionary model across both terrestrial carnivorans and pinnipeds (full dataset).

Figure S4. Clade-specific evolutionary shifts in skeletal components across both terrestrial carnivorans and pinnipeds (full dataset).

Figure S5. Phylomorphospaces of skeletal components and full phenome across both terrestrial carnivorans and pinnipeds (full dataset).

Figure S6. Clade-specific evolutionary shifts in the full phenome across both terrestrial carnivorans and pinnipeds (full dataset).

Supplementary Table Headings (full Supplementary Tables S1–S8 are in a separate .xlsx file)

References

**Supplementary Methods**

*Skeletal and ecological traits*

We used 103 measurements to capture the skeletal morphology of 208 osteological specimens across 119 extant species from 14 of the 16 carnivoran families (see Table S1 for list of specimens and museums). We were unable to collect data from Prionodontidae, which contains two extant species of Asiatic linsangs, and Odobenidae, which contains a single extant species (the walrus). Although we were able to gather data from only a third of carnivoran species diversity, our sample represents all major carnivoran ecomorphs. In total, we measured seven traits in the cranium; seven traits in the mandible; 13 traits in the forelimb (scapula, humerus, ulna, radius, third metacarpal); 13 traits in the hindlimb (pelvis, femur, tibia, fibula, calcarean, and third metatarsal); and seven traits in third cervical, fifth cervical, first thoracic, middle thoracic (the sixth thoracic if the species has 13 or 14 thoracic vertebrae or the seventh thoracic if the species has 15 or 16 thoracic vertebrae), diaphragmatic thoracic, last thoracic, first lumbar, middle lumbar (the third lumbar if the species has four, five, or six lumbar vertebrae or the fourth lumbar if the species has seven lumbar vertebrae), and last lumbar vertebrae (Fig. S1). These measurements were chosen based on previous investigations of distinct ecomorphology among carnivorans [1–5]. All measurements were taken using Mitutoyo digital calipers or a tape measure. All specimens were fully mature, determined by the closure of exoccipital-basioccipital and basisphenoid-basioccipital sutures on the cranium, full tooth eruption, and closure of sutures on the limb bones. We used only male specimens because carnivorans exhibit differing degree of sexual dimorphism [6,7].

To remove the effects of size, we calculated log shape ratios by dividing each skeletal trait by the geometric mean of all 103 traits as our metric of size (i.e., ln(trait/geometric mean) [8,9]. Species means were calculated prior to statistical analyses. We then used individual principal component analyses (PCAs) to reduce the dimension of each skeletal component (i.e., cranium, mandible, forelimb, hindlimb, and each of the nine vertebrae) and retained a number of PC axes that correspond to at least 90% of the explained variance. We then conducted a PCA on the entire dataset as our proxy of the overall skeletal body plan and retained the first six PC axes (>75% of explained variance) for subsequent analyses.

We classified the 119 carnivoran species into one of six locomotor categories (i.e., arboreal, semiarboreal, aquatic, semi-aquatic, semi-fossorial, and terrestrial), one of six hunting behavior categories (ambush, pounce, pursuit, occasional, semi-fossorial, and aquatic), and one of six dietary categories (carnivory, omnivory, insectivory, aquatic carnivory, and herbivory) following [10].

*Phylogenetic comparative methods*

We tested whether each skeletal component evolved as adaptations to different on diets, hunting behaviors, or locomotor modes or exhibited clade-based evolutionary shifts by fitting multivariate generalized evolutionary models on the retained PC axes of each skeletal component and the overall skeletal body plan [11–13]. For the adaptive ecological models, we three fit multi-optima Ornstein-Uhlenbeck models (i.e., mvOUM_diet_, mvOUM_hunting behavior,_ and mvOUM_locomotion_) to test if dietary, hunting behavioral, or locomotor regimes influenced the evolution of each set of PC axes using the R package mvMORPH version 1.1.4 [13]. These models allowed each ecological regimes to exhibit different trait optima. All three models were fit across 500 stochastically mapped trees to account for uncertainty in phylogenetic topology and the ancestral character states. We inferred the evolution of dietary, hunting behavioral, or locomotor regimes by performing stochastic character mapping with symmetric transition rates between regimes [14–16] in phytools [17]. We simulated 10 stochastic character maps across 1000 tree topologies randomly drawn from the posterior distribution of trees [18], resulting in 10,000-character maps for each set of diet, diet based on relative prey size, diet based on prey properties, and hunting behavior regimes. We randomly sampled 500 trees for subsequent analyses. We also calculated the phylogenetic half-lives (ln(2)/alpha) of the best supported adaptive ecological model, defined as the time it takes for a trait to evolve halfway toward its new expected optimum after a regime shift [11]. A short phylogenetic half-life relative to the age of Carnivora (48.2 myr) would suggest that skeletal traits are strongly pulled toward distinct ecological peaks across the adaptive landscape. For the clade-based model, we fit a multi-optima OU model (mvOUM_phyloEM_) without a priori ecological groupings with the R package PhylogeneticEM version 1.4.0 [19]. This data-driven approach can detect evolutionary shifts toward different optima without influences of a priori groupings on the tree. We used a scalar OU model that infers the full evolutionary rate matrix and accounts for correlations within multivariate datasets (i.e., PC axes). We also fit a single-rate multivariate Brownian motion model (mvBM1), which assumes trait variance accumulates stochastically but proportionally to evolutionary time, and a single-optimum Ornstein-Uhlenbeck model (mvOU1), which constrains each PC to evolve toward a single optimum. Relative support for each of the six models was assessed through computation of small sample-corrected Akaike weights (AICcW).

We acknowledge that using a subset of PC axes instead of the full dataset may lead to inaccurate results [20], but we are computationally limited to run evolutionary models on all traits or PC axes within each component. Nevertheless, we conducted a sensitivity analysis to determine if adding additional PC axes to our models changed our results. We added one more PC axis to each skeletal component and reran the evolutionary models in mvMORPH. Due to computational limits, we fit models across only 50 stochastically mapped trees instead of the full 500 trees. We also reran the clade-based model in PhylogeneticEM. We found that the addition of one PC axis to each component did not change our results except for the third cervical vertebrae modeling (Table S4). In this case, the best model was the single-optimum (OU1) model whereas the original best model was the clade-shift model. Despite the model difference, an adaptive ecological model was not the better fit and thus does not change the results of our manuscript.

Lastly, we assessed the covariation among skeletal components using two-block partial least squares (PLS) in the R package geomorph version 4.0.6 [21]. We conducted a PLS that assumes no phylogenetic structure with the geomorph function integration.test as well as a phylogenetic PLS that assume full Brownian motion with the geomorph function phylo.integration.

All statistical tests were performed on a reduced dataset that does not include the pinnipeds (results reported in the main text) and on a full dataset that does include the pinnipeds (results reported below).

**Supplementary Results and Discussion**

*Optima distribution in phylomorphospace*

The mvOUM_diet_ model was the best supported model for the mandible (Fig. 1; Table S2). PCs 1 and 2 explain 48.0% and 16.2% of the variance, respectively. Positive PC 1 scores are associated with relatively longer moment arms of the masseter and out-levers to the canine and molar; positive PC 2 scores are also associated with relatively longer moment arms of the masseter whereas negative PC 2 scores are associated with relatively longer out-levers to the canine and molar (Table S5). The mvOUM_diet_ model suggests that optima for herbivores and piscivores are at the extreme ends of PC 1 whereas optima for carnivores, omnivores, and insectivores are similar in phylomorphospace (Fig. S2B). These results are consistent with previous analyses in carnivorans using 3D geometric morphometrics [22].

The mvOUM_locomotion_ model was the best supported model for the hindlimb (Fig. 1; Table S2). PCs 1 and 2 explain 47.8% and 27.3% of the variance, respectively. Negative PC 1 scores are associated with relatively longer metatarsals, fibula, and tibia; positive PC 2 scores are associated with relatively broader greater trochanter of the femur (Table S5). The mvOUM_locomotion_ model suggests that optima for semi-fossorial carnivorans followed by semi-aquatic carnivorans are associated with positive PC 1 and negative 2 scores whereas optima for arboreal, semi-arboreal, and terrestrial carnivorans occupy a similar central region in phylomorphospace (Fig. S2D). Relatively shorter hindlimbs are beneficial semi-fossorial and semi-aquatic carnivorans: relatively shorter hindlimbs in semi-fossorial carnivorans can stabilize the torso against the large forces generated during digging with the forelimbs [23,24], and relatively shorter hindlimbs in semi-aquatic carnivorans bring the paddling limb closer to the body, and thus reduces induced drag during the recovery stroke [4,25,26].

The mvOUM_locomotion_ model was also the best supported model for the middle lumbar vertebrae (Fig. 1; Table S2). PCs 1 and 2 explain 59.1% and 16.2% of the variance, respectively. Positive PC 1 scores are associated with overall vertebral bodies that are relatively wider and taller and centra that are relatively taller in height but shorter in length; negative PC 2 scores are associated with relatively taller neural spine and the overall vertebral body in height, centra that are relatively longer and taller, and relatively longer inter-zygapophyseal lengths (Table S5). The mvOUM_locomotion_ model suggests that the optimum for arboreal carnivorans are associated with negative PC 1 and positive PC 2 scores whereas semi-aquatic carnivorans are associated with positive PC 1 and negative PC 2 scores (Fig. S2L). In contrast, optima for semi-fossorial, semi-arboreal, and terrestrial carnivorans occupy a similar central region in phylomorphospace (Fig. S2L).

The mvOUM_hunting behavior_ model was the best supported model for both the last thoracic, first lumbar, and last lumbar vertebrae (Fig. 1; Table S2). In the last thoracic vertebrae, PCs 1 and 2 explain 65.3% and 15.4% of the variance, respectively. Negative PC 1 scores are associated with relatively taller vertebrae including the neural spines and centra but relatively shorter lengths of the centra; negative PC 2 scores are associated with relatively shorter centra in both height and length and relatively longer inter-zygapophyseal lengths (Table S5). In the first lumbar vertebrae, PCs 1 and 2 explain 67.2% and 12.7% of the variance, respectively. Negative PC 1 scores are associated with relatively taller neural spines and vertebral that are both relatively taller and wider; negative PC 2 scores are associated with relatively shorter centra in both height and length and relatively longer inter-zygapophyseal lengths (Table S5). In the last lumbar vertebrae, PCs 1 and 2 explain 47.9% and 29.4% of the variance, respectively. Negative PC 1 scores are associated with relatively taller vertebrae including the neural spines and centra; negative PC 2 scores are associated with relatively longer centrum lengths and relatively longer inter-zygapophyseal lengths (Table S5). The optima distributions were similar in the last thoracic and first lumbar vertebrae (Fig. S2J, K). Pursuit carnivorans followed by aquatic and semi-fossorial carnivorans are associated with negative PC 1 scores, whereas optima for ambush and occasional carnivorans occupy a similar central region in phylomorphospace (Fig. S2J, K). The optima for pursuit, aquatic, and semi-fossorial carnivorans exceed the boundaries of extant species in phylomorphospace, suggesting that these optima are poorly estimated. A possible reason for these poor estimations is the low number of species in the pursuit (n = 5), semi-fossorial (n = 5), and aquatic (n = 7) regimes. In addition, uncertainty in optimal estimations can also arise when species in the regime are spread across the phylogeny. That the pursuit and semi-fossorial carnivorans are spread across the entire carnivoran phylogeny and share a common ancestor over 48 million years ago may contribute to high uncertainty in optimal estimates. The optima distribution in the last lumbar is slightly different compared to the last thoracic and first lumbar vertebrae (Fig. S2M). Except for pouncing carnivorans, the optima for all other hunting behavioral regimes are associated with negative PC 1 scores. Pursuit carnivorans are also associated with positive PC 2 scores, where shorter vertebral lengths may increase flexibility when chasing after prey.

*Patterns of covariation*

All integration tests among skeletal components were statistically significant except for the covariation between the hindlimb and transitional thoracic vertebrae (Table S3). The strongest covariations occur within the skull (i.e., cranium and mandible), appendicular system (i.e., forelimb and hindlimb), and among the vertebral regions of the axial skeleton (i.e., third and fifth cervical vertebrae, among anterior thoracic vertebrae, and among posterior thoracic and lumbar vertebrae) (Table S3). Skeletal components that share the same evolutionary model do not necessarily exhibit stronger covariation. For example, the OUM_locomotion_ model was the best fitting model for the hindlimb and middle lumbar vertebrae (Table S2); however, the correlation coefficient between these two skeletal components is similar as the correlation coefficients between the hindlimb and other thoracic and lumbar vertebrae (Table S3). The distribution of locomotor optima varies between the hindlimb and middle lumbar (Fig. S2D, L), which may decrease their evolutionary covariation even though the same extrinsic factor influences their evolution. Another possible explanation is that the OUM_hunting_ model was a similarly good fit for the middle lumbar vertebrae (ΔAICc = 1.19; Table S2), which is in line with the last thoracic, first lumbar, and last lumbar vertebrae. These vertebrae exhibited high correlation coefficients (Table S3). Overall, our covariation analyses are consistent with previous work finding that elements within and among the skull, appendicular, and axial skeletal systems typically show significant covariation that may weaken or strengthen from extrinsic factors (e.g., [24,27–34]).

*Analyses with full dataset*

Like analyses with just the terrestrial carnivorans, we found that the clade-specific shift model exhibited overwhelmingly greater support in the cranium (mvOUM_phyloEM_; AICcW > 0.99) whereas the adaptive ecological model based on diet was the best supported model (mvOUM_diet_; AICcW = 0.96) for the mandible (Fig. S3; Fig. S5B; Table S6). However, we found only two evolutionary shifts in cranial morphology occurring on the branches towards Arctoidea and pinnipeds (Fig. S4A).

In the appendicular skeleton, the clade-specific shift model was the best supported for both forelimb and hindlimb (Fig. S3; Table S6). The forelimb exhibited similar evolutionary shifts as in the reduced analyses with the addition of two shifts corresponding to each of the two represented pinniped clades: Phocidae and Otariidae (Fig. S4B). The hindlimb exhibited a single evolutionary shift that corresponded to the pinniped clade (Fig. S4C); excluding the clade-specific shift model revealed that adaptive ecological model based on locomotion was the best model (Table S6), the same result when analyzing the reduced dataset with no pinnipeds.

In the axial skeleton, there were no distinct modes of evolution between the anterior and posterior regions of the vertebrae column as found in the analyses with the reduced dataset with no pinnipeds. Instead, the clade-specific shift model was the best supported model for all sampled vertebrae (Fig. S3; Table S6). However, the last thoracic, first lumbar, and middle lumbar vertebrae all exhibited a single evolutionary shift towards or within the pinniped clade (Fig. S4I–K). This suggests that the divide between terrestrial carnivorans and pinnipeds are the major source of variation in these vertebral morphologies, and excluding the clade-specific shift model revealed that adaptive ecological model based on locomotion was the best model for all three vertebrae (Table S6).

The clade-specific shift model was the best supported model for the full phenome dataset (Fig. S6; Table S6). We found the same seven evolutionary shifts in the full phenome dataset that correspond to carnivoran clades with the addition of two shifts corresponding to each of the two represented pinniped clades: Phocidae and Otariidae (Fig. S5).

All integration tests among skeletal components were statistically significant (Table S7). Correlation coefficients are similar to the reduced dataset (Table S3).

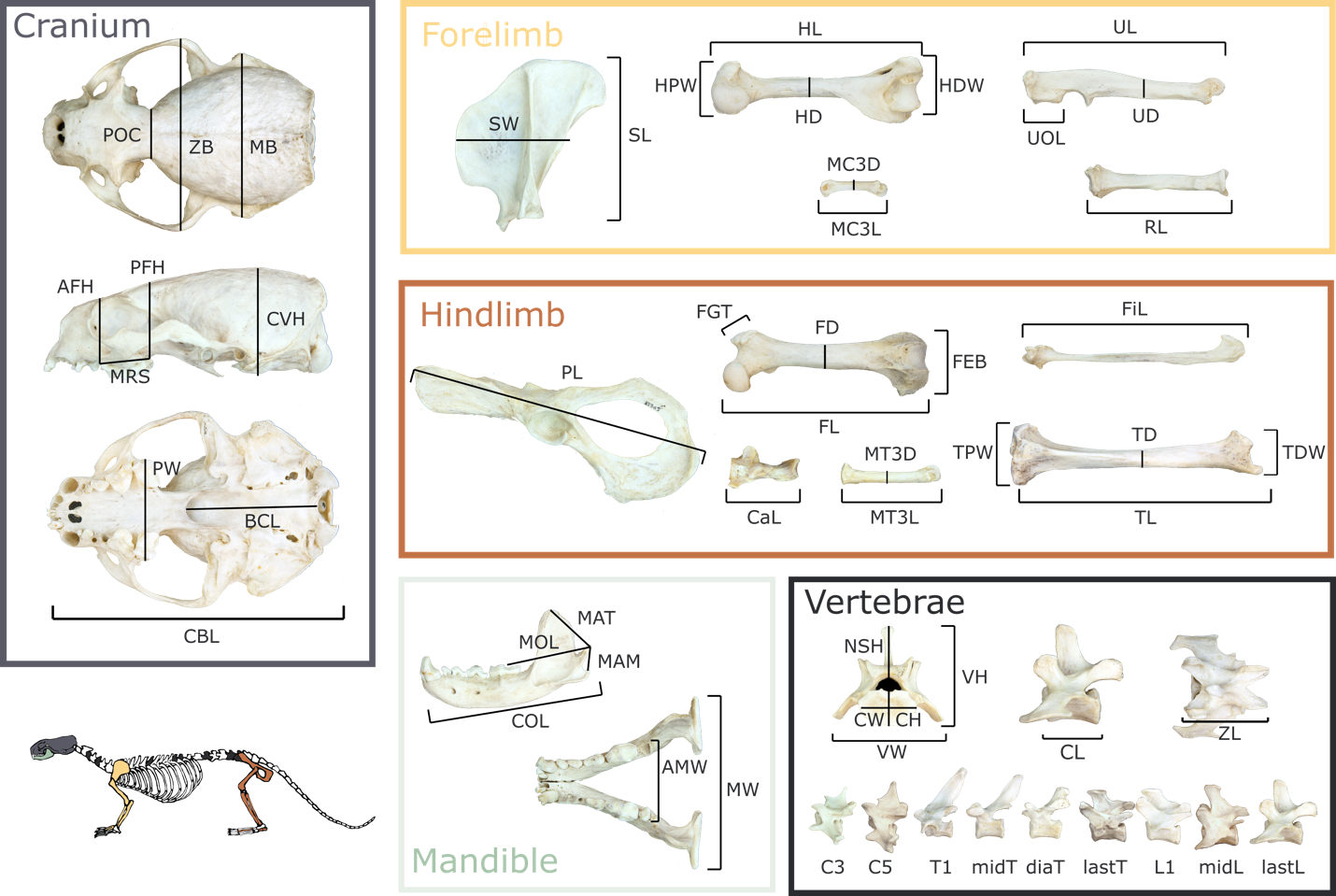

**Fig. S1.** Skeletal trait measurements used in this study. *Cranium*: CBL = condylobasal length, POC = postorbital constriction breadth, ZB = zygomatic breadth, MB = mastoid breadth, AFH = anterior facial height, PFH = posterior facial height, CVH = cranial vault height, PW = palate width, BCL = basicranial length. *Mandible*: MAT = moment arm of temporalis, MAM = moment arm of masseter, MAM2 = moment arm of masseter; MOL = molar out-lever; COL = canine out-lever; MW = mandibular width; AMW = anterior mandibular width. *Forelimb*: SL = scapula length; SW = scapula width; HL = humerus length; HD = humerus mid-shaft width; HPW = humerus proximal width; HDW = humerus distal width; UL = ulna length; UD = ulna mid-shaft width; UOL = ulnar olecranon length; RL = radius length; RD = radius mid-shaft width; MC3L = third metacarpel length; MC3W = third metacarpel width. *Hindlimb*: PL = pelvis length; FL = femur length; FD = femur mid-shaft width; FEB = femur distal width; FGT = height of the greater trochanter of the femur; TL = tibia length; TD = tibia mid-shaft width; TPW = tibia proximal width; TDW = tibia distal width; FiL = fibula length; CaL = calcaneus length; MT3L = third metatarsal length; MT3D = third metatarsal width. *Vertebrae*: VW = vertebrae width; VH = vertebrae height; CL = centrum length; CW = centrum width (posterior); CH = centrum height (posterior); ZL = inter-zygapophyseal length; NSH = neural spine height.

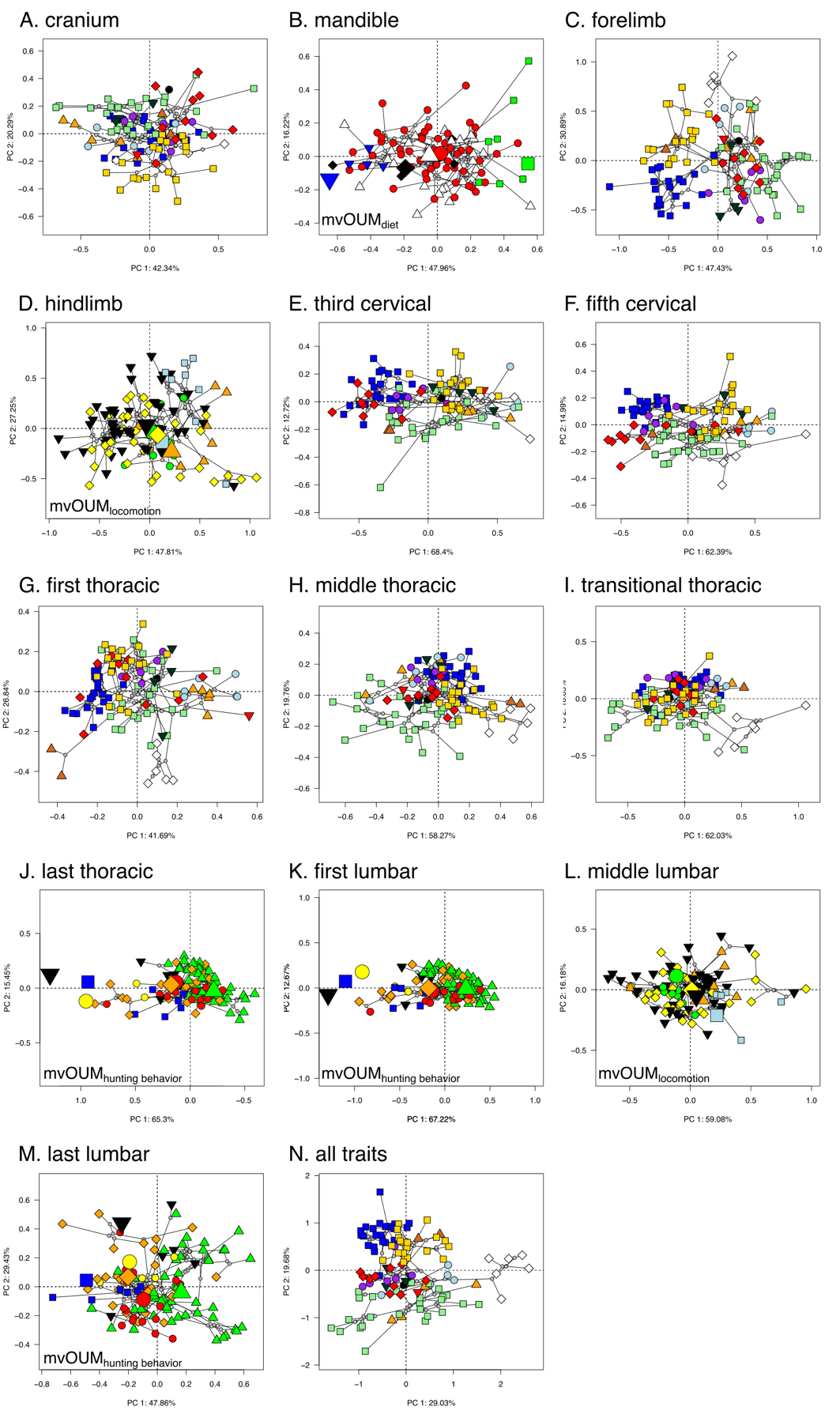

**Fig. S2.** Phylomorphospace of (A–M) skeletal components and (N) full phenome across terrestrial carnivorans. Species tips in the cranium; forelimb; cervical vertebrae; and first, middle, and transitional vertebrae are color coded based on family names: black circle = Nandiniidae; gold square = Felidae; red diamond = Viverridae; brown triangle = Hyaenidae; dark green upside down triangle = Eupleridae; purple circle = Herpestidae; blue square = Canidae; white diamond = Ursidae; orange triangle = Mephitidae; red upside down triangle = Ailuridae; light blue circle = Procyonidae; light green square = Mustelidae. Species tips in the mandible are color coded based on dietary regimes: red circle = carnivory; green square = herbivory; black diamond = insectivory; white triangle = omnivory; blue upside down triangle = piscivory. Species tips in the hindlimb and middle lumbar vertebrae are color coded based on locomotor regimes: green circle = arboreal; light blue square = semi-aquatic; yellow diamond = semi-arboreal; orange triangle = semi-fossorial; black upside down triangle = terrestrial. Species tips in the last thoracic, first lumbar, and last lumbar vertebrae are color coded based on hunting behavioral regimes: red circle = ambush; blue square = aquatic; orange diamond = occasional; green triangle = pounce; black upside down triangle = pursuit; yellow circle = semi-fossorial. Larger symbols represent the optimum of each respective regime estimated from the best fitting evolutionary model. ­

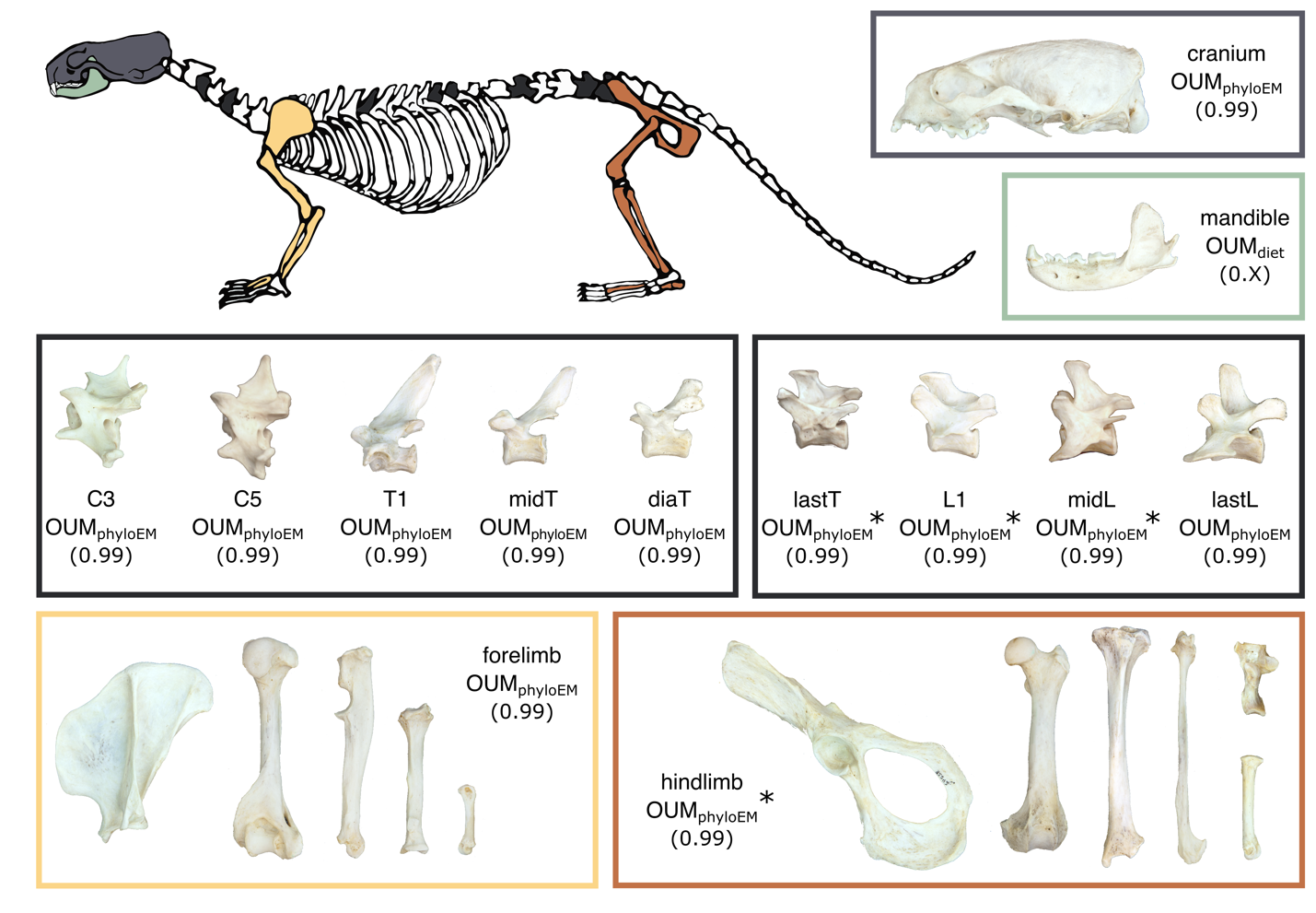

**Fig. S3.** Diagram of the skeletal components and their best-fitting evolutionary model across both terrestrial carnivorans and pinnipeds. AICcW are in parentheses. See Table S6 for full AICc table. C3 = third cervical vertebrae; C5 = fifth cervical vertebrae; T1 = first thoracic vertebrae; midT = middle thoracic vertebrae; diaT = diaphragmatic thoracic vertebrae; lastT = last thoracic vertebrae; L1 = first lumbar vertebrae; midL = middle lumbar vertebrae; lastL = last lumbar vertebrae. *indicates that there is only a single evolutionary shift within the OUM_phyloEM_ model that corresponds to pinnipeds (Fig. S4); in all cases, the OUM_locomotion_ model is the second best model.

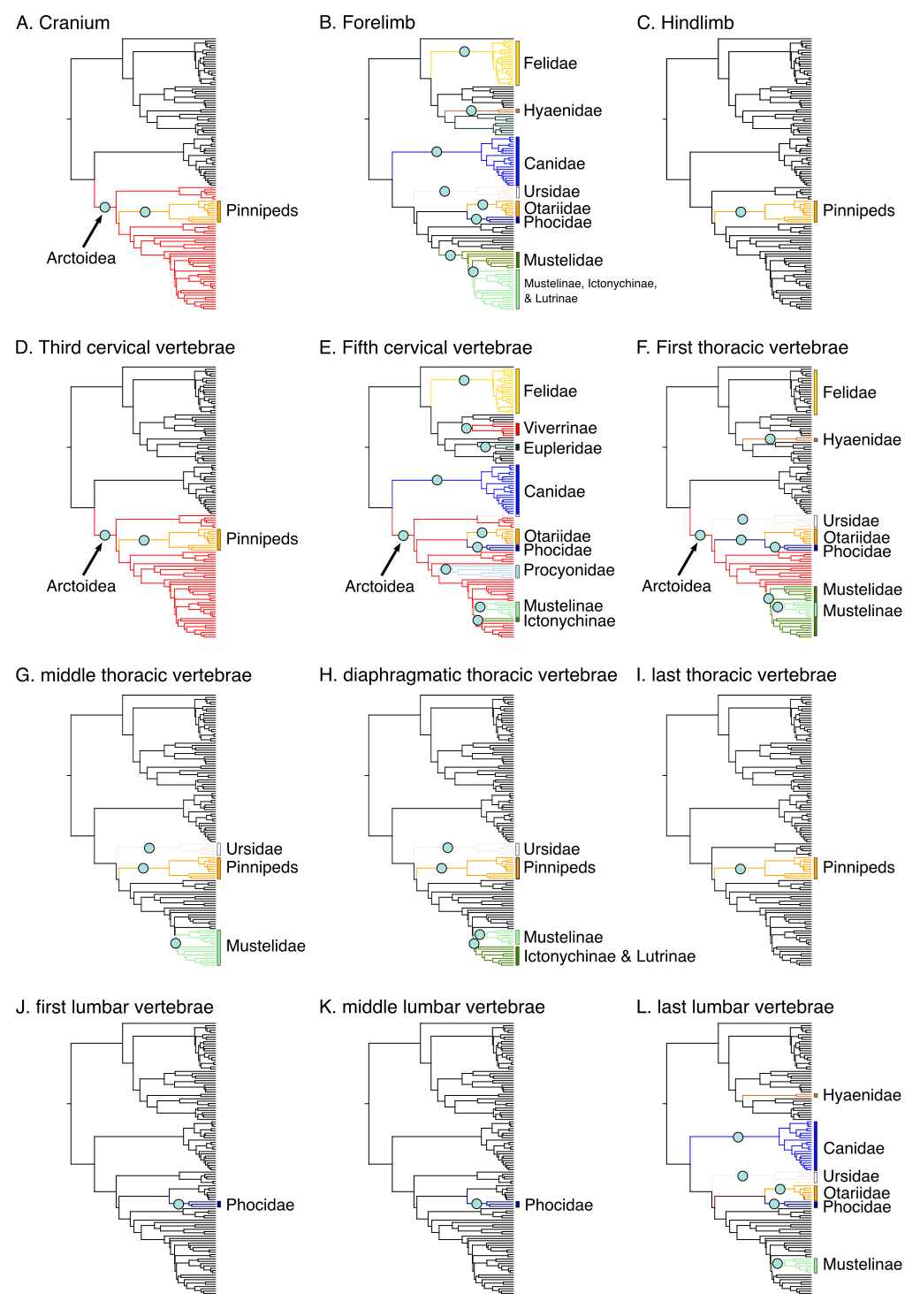

**Fig. S4.** Clade-specific evolutionary shifts in skeletal components across both terrestrial carnivorans and pinnipeds identified by PhylogeneticEM. Shifts are represented as pink circles, and branches on the phylogenies are colored according to each regime.

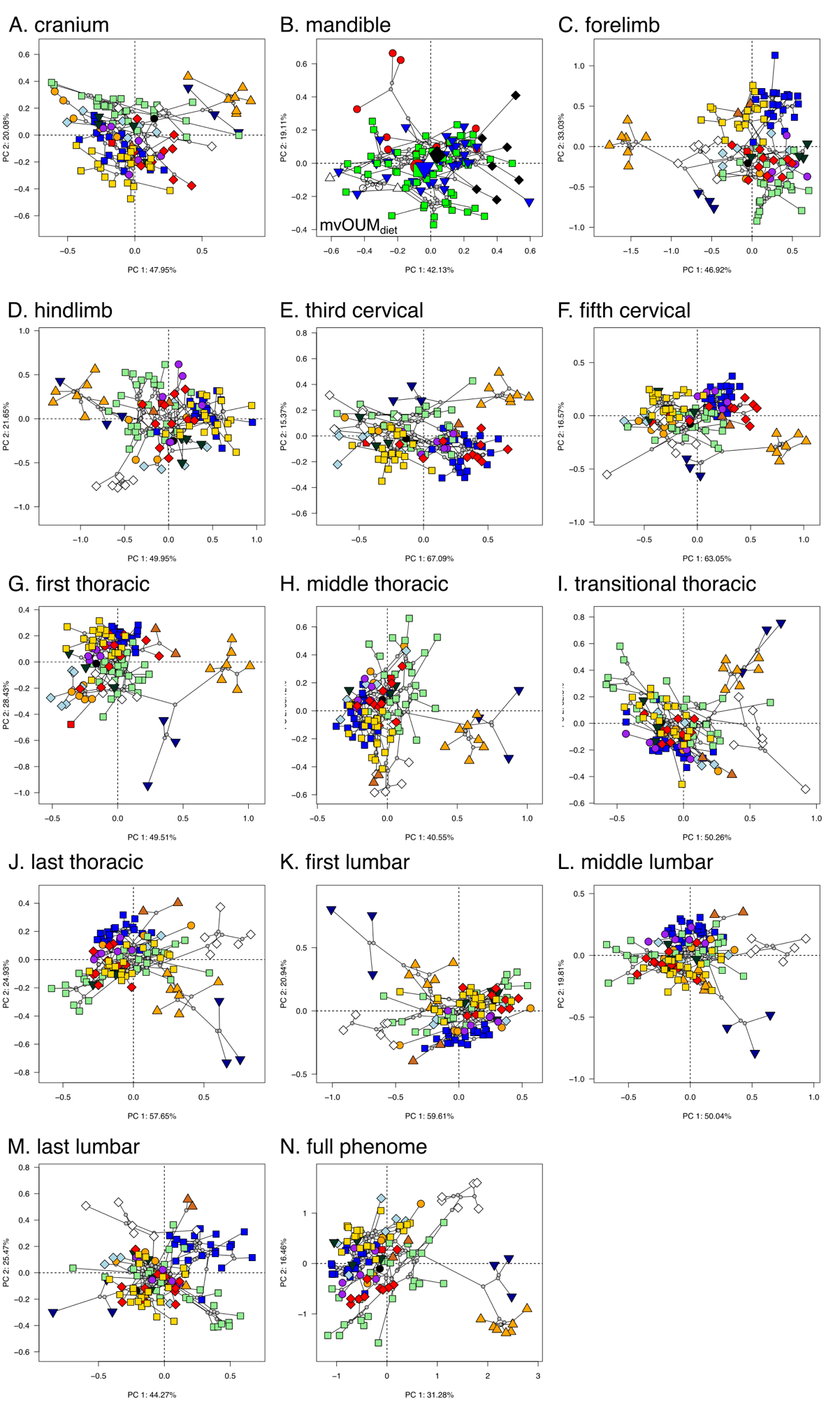

**Fig. S5.** Phylomorphospace of (A–M) skeletal components and (N) full phenome across terrestrial carnivorans and pinnipeds (full dataset). Species tips all components except the mandible are color coded based on family names: black circle = Nandiniidae; gold square = Felidae; red diamond = Viverridae; brown triangle = Hyaenidae; dark green upside down triangle = Eupleridae; purple circle = Herpestidae; blue square = Canidae; white diamond = Ursidae; orange triangle = Mephitidae; red upside down triangle = Ailuridae; light blue circle = Procyonidae; light green square = Mustelidae. Species tips in the mandible are color coded based on dietary regimes: red circle = carnivory; green square = herbivory; black diamond = insectivory; white triangle = omnivory; blue upside down triangle = piscivory. Larger symbols represent the optimum of each respective regime estimated from the best fitting evolutionary model. ­

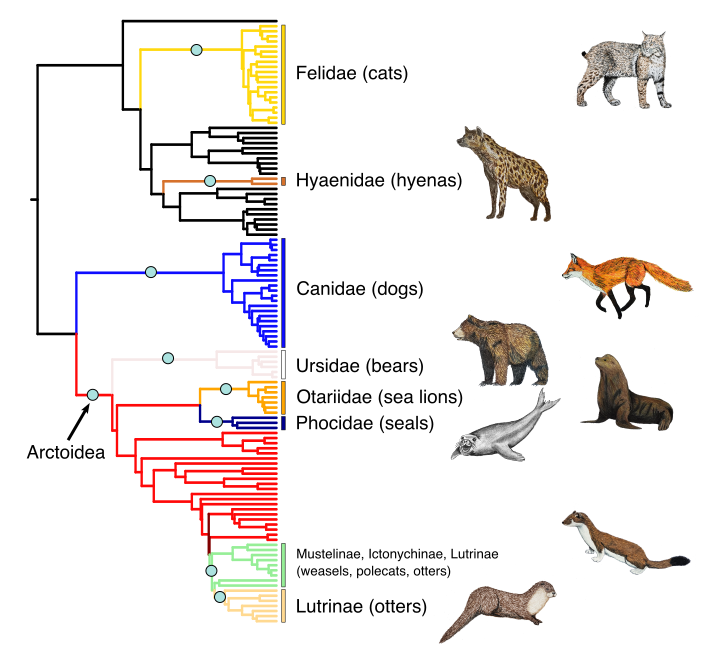

**Fig. S6.** Clade-specific evolutionary shifts of the full skeletal dataset across terrestrial carnivorans and pinnipeds (full dataset) identified by PhylogeneticEM. Shifts are calculated from PCs 1-6 (77.8% of explained variance) and are represented as pink circles. Branches on the phylogenies are colored according to each regime.

**Table headings**

**Table S1.** Specimens and museum catalog numbers used in this study. AMNH = American Museum of Natural History; CAS = California Academy of Sciences; FMNH = Field Museum of Natural History; LACM = Natural History Museum of Los Angeles County; MVZ = Museum of Vertebrate Zoology; NHMUK = Natural History Museum, London; SDNHM = San Diego Natural History Museum; TxVP = Texas Vertebrate Paleontology Collection; USNM = National Museum of Natural History; UWBM = Burke Museum of Natural History and Culture

**Table S2.** Comparisons of the best-fitting evolutionary models in each of the skeletal components in terrestrial carnivorans (reduced dataset). Small sample-corrected Akaike weights (AICcW) were calculated for each of the 500 replications to account for uncertainty in phylogenetic topology and the ancestral character states. Rows in boldface type represent the best-fit model as indicated by the lowest ΔAICc score. ΔAICc = the mean of AICc minus the minimum AICc between models.

**Table S3**. Results of two-block partial least squares (PLS) among skeletal components across terrestrial carnivorans (reduced dataset). Numbers represent correlation coefficient (r). Bolded numbers indicate P < 0.05.

**Table S4.** Sensitivity test of adding additional PC axis to each component. Comparisons of the best-fitting evolutionary models in each of the skeletal components in terrestrial carnivorans (reduced dataset). Small sample-corrected Akaike weights (AICcW) were calculated for each of the 50 replications to account for uncertainty in phylogenetic topology and the ancestral character states. Rows in boldface type represent the best-fit model as indicated by the lowest ΔAICc score. ΔAICc = the mean of AICc minus the minimum AICc between models.

**Table S5**. Trait loadings from PCAs of each component across terrestrial carnivorans (reduced dataset). Trait abbreviations are defined in Fig. S1.

**Table S6.** Comparisons of the best-fitting evolutionary models in each of the skeletal components across both terrestrial carnivorans and pinnipeds (full dataset). Small sample-corrected Akaike weights (AICcW) were calculated for each of the 500 replications to account for uncertainty in phylogenetic topology and the ancestral character states. Rows in boldface type represent the best-fit model as indicated by the lowest ΔAICc score. ΔAICc = the mean of AICc minus the minimum AICc between models.

**Table S7**. Results of two-block partial least squares (PLS) among skeletal components across terrestrial carnivorans and pinnipeds (full dataset). Numbers represent correlation coefficient (r). Bolded numbers indicate P < 0.05.

**Table S8**. Trait loadings from PCAs of each component across terrestrial carnivorans and pinnipeds (full dataset). Trait abbreviations are defined in Fig. S1.
